# High-speed multifocal plane fluorescence microscopy for three-dimensional visualisation of beating flagella

**DOI:** 10.1101/573485

**Authors:** Richard J. Wheeler

## Abstract

Analysis of flagellum beating in three dimensions is important for understanding how cells can undergo complex flagellum-driven motility and the ability to use fluorescence microscopy for such three-dimensional analysis would be extremely powerful. *Trypanosoma* and *Leishmania* are unicellular parasites which undergo complex cell movements in three dimensions as they swim and would particularly benefit from such an analysis. Here, high-speed multifocal plane fluorescence microscopy, a technique in which a light path multi-splitter is used to visualise 4 focal planes simultaneously, was used to reconstruct the flagellum beating of *Trypanosoma brucei* and *Leishmania mexicana* in three dimensions. It was possible to use either an organic fluorescent stain or a genetically-encoded fluorescence fusion protein to visualise flagellum and cell movement in three dimensions at a 200 Hz frame rate. This high-speed multifocal plane fluorescence microscopy approach was used to address two open questions regarding *Trypanosoma* and *Leishmania* swimming: To quantify the planarity of the *L. mexicana* flagellum beat and analyse the nature of flagellum beating during *T. brucei* ‘tumbling’.

## Introduction

Trypoanosomatid parasites, including the human pathogens *Trypanosoma brucei*, *Trypanosoma cruzi* and *Leishmania* spp., have a single flagellum whose motility is vital for progression through the life cycle. They are genetically tractable and an excellent model for flagellum biology. Different species and life cycle stages have differences in motility linked with morphological adaptations and these morphologies are defined, in part, by the flagellum. The flagellum is laterally attached to the cell body along most of its length in the trypomastigote morphology, for a shorter portion of its length in the epimastigote morphology and is not laterally attached to the cell body (instead protruding from the cell anterior) in the promastigote morphology^1^.

*Trypanosoma brucei* trypomastigotes, including the procyclic (fly midgut) and mammalian bloodstream forms, have a complex three-dimensional (3D) cell movement^2–5^ giving rise to motility vital for the parasite^6^. They can swim in liquid, but are not always free in a large fluid volume^3^, and when they are they may be swimming at several orders of magnitude slower than the fluid flow^5,7^. Furthermore, they can invade tissues^8^. Together, this speaks to multiple functions of flagellum motility, now known to include complex collective motion of densely packed cells^9–11^ and surface antibody clearance^12,13^. The motility of *Trypanosoma cruzi* and *Leishmania* have been less extensively analysed, however flagellum motility is likely similarly multifunctional^14^. Swimming of *Leishmania* promastigotes and *T. cruzi* epimastigotes have also been analysed and cell movement and swimming behaviours are somewhat unlike *T. brucei*^2,15,16^. As for *T. brucei*, they are not always free in a large fluid volume and, when densely packed, will also likely undergo hydrodynamic coupling of movement which could give collective movement of a population at the macro scale.

Contributing to complex motility are at least two modes of flagellum movement. *T. brucei*, *T. cruzi, Leishmania* and related organisms normally undergo a tip to base symmetrical (flagellar type) beat. In *Leishmania* promastigotes and *T. cruzi* epimastigotes this is near-planar^2,16,17^. These organisms can also switch to a base to tip beat^5,15–18^. In *T. cruzi* epimastigotes and *Leishmania* promastigotes this reversed beat is asymmetrical (a ciliary type beat) and causes a rotation of cell orientation^15,16,18^. In *T. brucei* the reversed beat is not so well characterised (outside of mutants with a reversed beat^19^) but occurs at a lower frequency and with a higher amplitude^5^ and can cause slow backwards swimming and ‘tumbling’^5^ - rapid cell reorientation associated with a contorted cell shape^20^.

Existing analysis of trypanosomatid cell/flagellum movement is two-dimensional with three notable exceptions for *T. brucei*^2,5^. One used defocus of bright-field microscopy to qualitatively infer distance from the focal plane^5^. Quantitative efforts were an optical tomography approach^5^ and mathematical model fitting^2^, however these both required a perfectly repeating beat and a constant rate of cell rotation while swimming. More broadly, three main approaches have been used to perform 3D analysis of beating: 1) Inferring defocus distance using bright-field microscopy^21^, similar to as previously used for *T. brucei*^5^. 2) Using a high frequency oscillating objective (essentially rapid focus change) to visualise multiple focal planes near-synchronously^22^. 3) High-speed digital holographic microscopy, used very effectively for the *Plasmodium* microgamete^23^ and sperm^24^. These approaches have been used ‘label free’, using transmitted light illumination.

Multifocal plane fluorescence microscopy^25,26^ has the potential to achieve the necessary frame rates and sensitivity to analyse flagellum beating using fluorescence in 3D with the advantage of using specific fluorescent labels. Here, high-speed multifocal plane fluorescence microscopy was used on *T. brucei* and *Leishmania mexicana* to analyse open questions about their flagellum beating. Firstly, signal from a membrane stain, FM 4-64FX, was used to analyse *T. brucei* flagellum beating during tumbling. This showed tumbling arises from flagellum motion which is not a simple time reversal of the normal flagellum beat with the proximal flagellum ‘locked’ in a curved configuration. Secondly, signal from a fluorescent protein, mNeonGreen (mNG)^27^, fused to a flagellar membrane protein at the endogenous locus was used to quantify the planarity of the *L. mexicana* flagellar beat. This showed flagellum bending remains within a plane but that plane can change over a beat cycle.

## Results and Discussion

Multifocal plane imaging was initially tested with *T. brucei* labelled with a fluorescent membrane stain, FM4-64, in normal growth medium. *T. brucei* movement deviates significantly from a single plane, with the flagellum beating is at around 20 Hz, requiring 100 Hz or faster frame rates to effectively ‘freeze’ flagellum/cell movement for analysis. Multifocal plane imaging was configured using a commercially available light path multi-splitter with three semi-silvered mirrors and lenses at the pupil planes to split the light to four quadrants of a single camera with focus offsets (Figure S1A). The lens focal lengths were selected for a focal plane (z) spacing of ~2 µm and the actual focus offset and the offset of sub-image position on the camera sensor (in x and y) were calibrated using fluorescent beads (Figure S1). Sub-image image offset and scale/magnification aberration was measured and corrected (Figure S1B, C, D) allowing precise alignment of the sub-images into a final z (focus) stack.

A 200 Hz frame rate was achieved with an image resolution of 2560 by 1080 px. At four focal planes this corresponds to ~0.5 gigavoxel/second with around ~8 photons/px background and signal up to ~50 photons/px (Figure 1A). Alignment of the sub-images in each quadrant and colour coding by focal plane offset gives a 3D view of cell conformation (Figure 1B) and image filtering highlights the in-focus membrane at each focal plane (Figure 1C). This effectively captured *T. brucei* morphology (Figure 1D); the contours of the cell and flagellum membrane can interpreted to give a plausible 3D configuration including the position of the flagellum and flagellar pocket at its base (Figure 1E). Alternatively, the data can be directly visualised in 3D (Figure 1F). This is the 3D configuration of the *T. brucei* cell at a well-resolved moment during flagellum movement.

**Figure 1.**
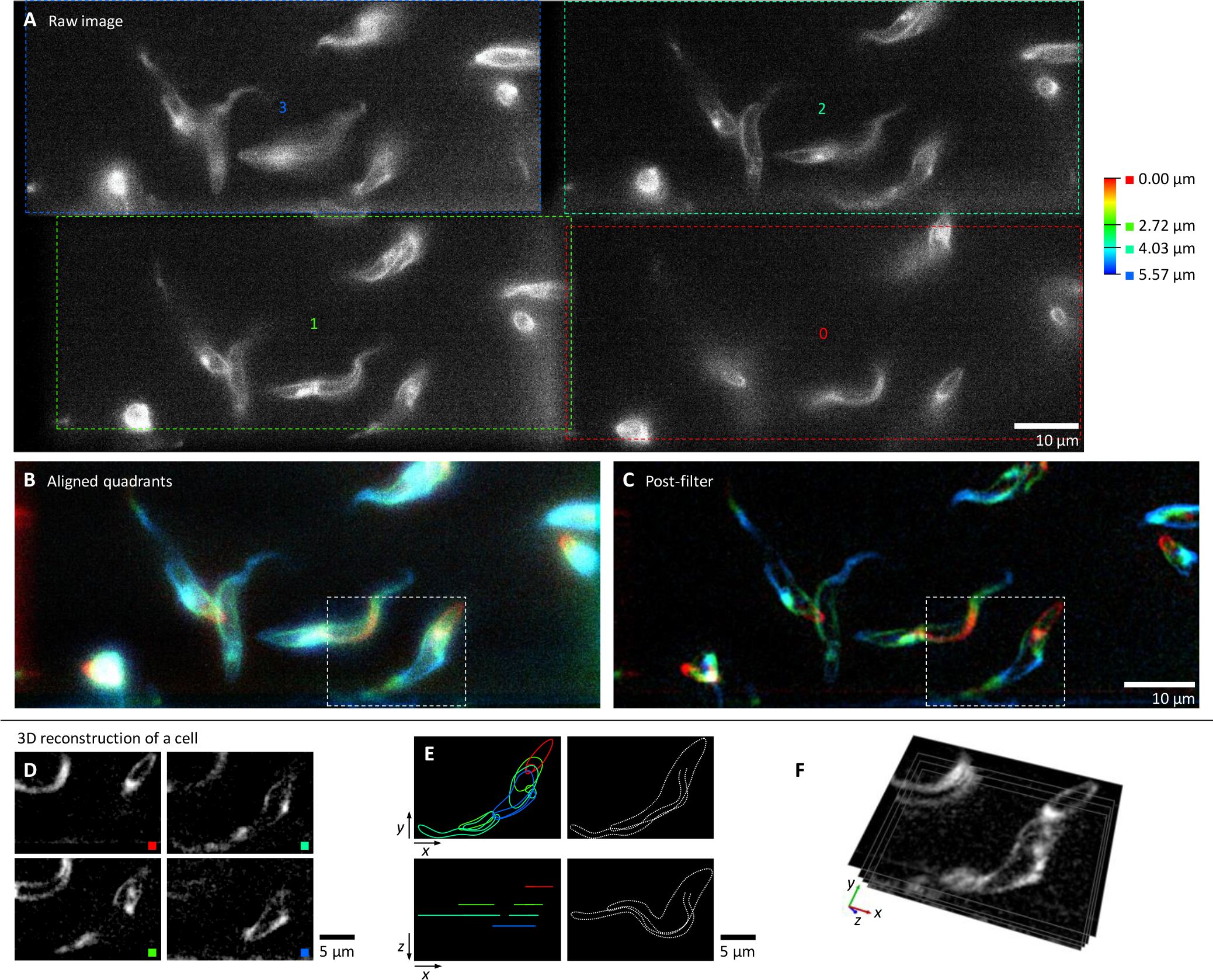
High-speed multifocal plane microscopy to visualise live swimming *T. brucei* in 3D. All panels show a single frame from a 200 Hz multifocal plane video of *T. brucei* procyclic forms labelled with a fluorescent membrane stain. **A.** The raw camera image, showing the four sub-images (0 to 3) and their focal offsets. **B.** The image in A, following alignment of the sub-images, pseudocoloured according to the depth scale in A. **C.** The image in B following filtering to aid interpretation. **D-F**. 3D reconstruction of an example cell, outlined in B and C. **D.** The individual filtered focal planes. **E.** Interpretation of the 3D conformation of the cell. Outlines from each focal plane are shown on the left and a sketch on the right. **F.** Direct visualisation of the focal planes in 3D.

### Flagellum beat during *T. brucei* tumbling

*T. brucei* procyclic and bloodstream form trypomastigotes can undergo at least two types of movement when in suspension culture, a directional forward swimming and a ‘tumbling’ associated with reversal of the beat. However, in the low Reynolds number environment *T. brucei* experience, a simple reversal of the flagellum beat with the same waveform would be expected to simply reverse swimming direction^28^. There may be an unrecognised asymmetry in the reversed *T. brucei* beat or perhaps there is some other change - a difference in motion between the proximal and distal flagellum has previously been noted^5^. Analysing this is challenging, as tumbling bloodstream form cells are associated with a ‘less straight’ configuration^21^ which sits poorly in a single focal plane.

To characterise flagellum movement while tumbling, samples of procyclic form cells were labelled with FM4-64 and placed in a sample chamber where tumbling cells sediment at the bottom (near the coverslip). 200 Hz multifocal plane videos of these cells showed all tumbling cells were undergoing a reversed (ie. base to tip) flagellum beat, however in the majority (75%, *n* = 28) the proximal half of the flagellum did not move and the waveforms instead initiated in the middle of the flagellum and propagated towards the tip (Figure 2A). In the remainder the entire flagellum moved and the waveforms initiated near the base of the flagellum (Figure 2B).

**Figure 2.**
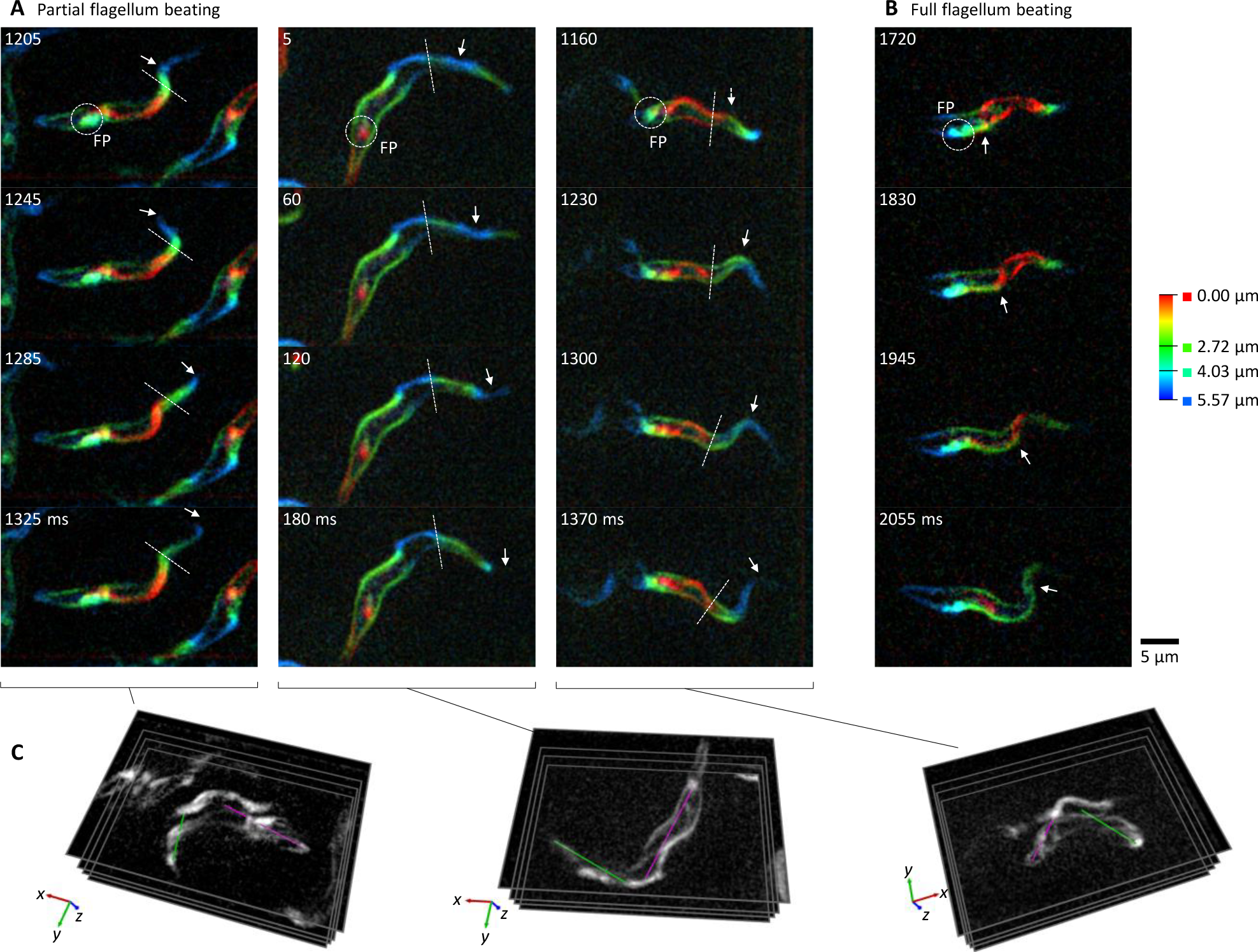
Tumbling *Trypanosoma brucei* tend to undergo a reversed flagellum beat in the distal half of the flagellum. All panels show frames from 200 Hz multifocal plane videos of *T. brucei* procyclic forms labelled with a fluorescent membrane stain. **A.** Three examples of the most common flagellum movement in ‘tumbling’ *T. brucei* procyclic cells showing four frames covering one beat cycle (beat frequency ~5 Hz). FP indicates the flagellar pocket at the base of the flagellum, arrows indicate a propagating wavefront, dotted line indicates the boundary between the beating and ‘locked’ portions of the flagellum. **B.** One example of the less common flagellum movement in tumbling cells where waves propagate along the entire flagellum. **C.** 3D conformation of the cells in A indicating the beating (green) and locked (magenta) portions of the flagellum, which lie approximately perpendicular.

In cells where flagellum movement was restricted to the distal half of the flagellum, the proximal flagellum was ‘locked’ in a curve on the time scale of 100 ms (Figure 2A). The 3D conformation of these cells (Figure 2C) showed the portion of the cell laterally attached to the beating part of the flagellum was oriented perpendicular to the portion of the cell not contorted by flagellar beating. The locked portion of the flagellum may explain the previously-described phenomenon of tumbling cells being less straight^21^. It also indicates a plausible origin of tumbling rather than directed swimming. Flagellum motion is not a simple time reversal of the normal tip to base beat and as it pushes the asymmetrically curved cell this leads to rotation.

For forward swimming waveforms can effectively initialise in the distal flagellum and propagate to the base, but, for tumbling, flagellar waveforms appear not to effectively initialise in the proximal half of the flagellum. Instead they initialise in the mid flagellum and then propagate to the tip. Propensity for waveform initiation at different sites may be related to differences in the molecular composition of outer dynein arms between the proximal and distal axoneme^19^. Alternatively, the higher stiffness of the wider cell body (which is laterally attached to the flagellum towards its base)^29^ may simply be too rigid for the reversed beat to bend.

### Planarity of the procyclic *L. mexicana* beat

The *Leishmania* flagellum beat is assumed to be planar, however *Leishmania* (and closely related species) flagellar beating has only previously been analysed when confined to a thin layer^2,17,18^. In addition, evidence for planarity is weak: it uses the criterion that the flagellum appears in focus along its entire length, for a beat which is oriented parallel to the focal plane. This is a qualitative assessment, limited by the z resolution of the microscope. To address this, *L. mexicana* promastigotes expressing an abundant flagellum membrane protein SMP1^30^ fused to mNG, SMP1::mNG, were analysed by multifocal plane microscopy. Videos were captured in a sample chamber in medium, focused away from the coverslip to avoid surface effects. A frame rate of 200 Hz could be achieved, with <1 photons/px background and signal up to 15 photons/px (Figure 3A,B).

**Figure 3.**
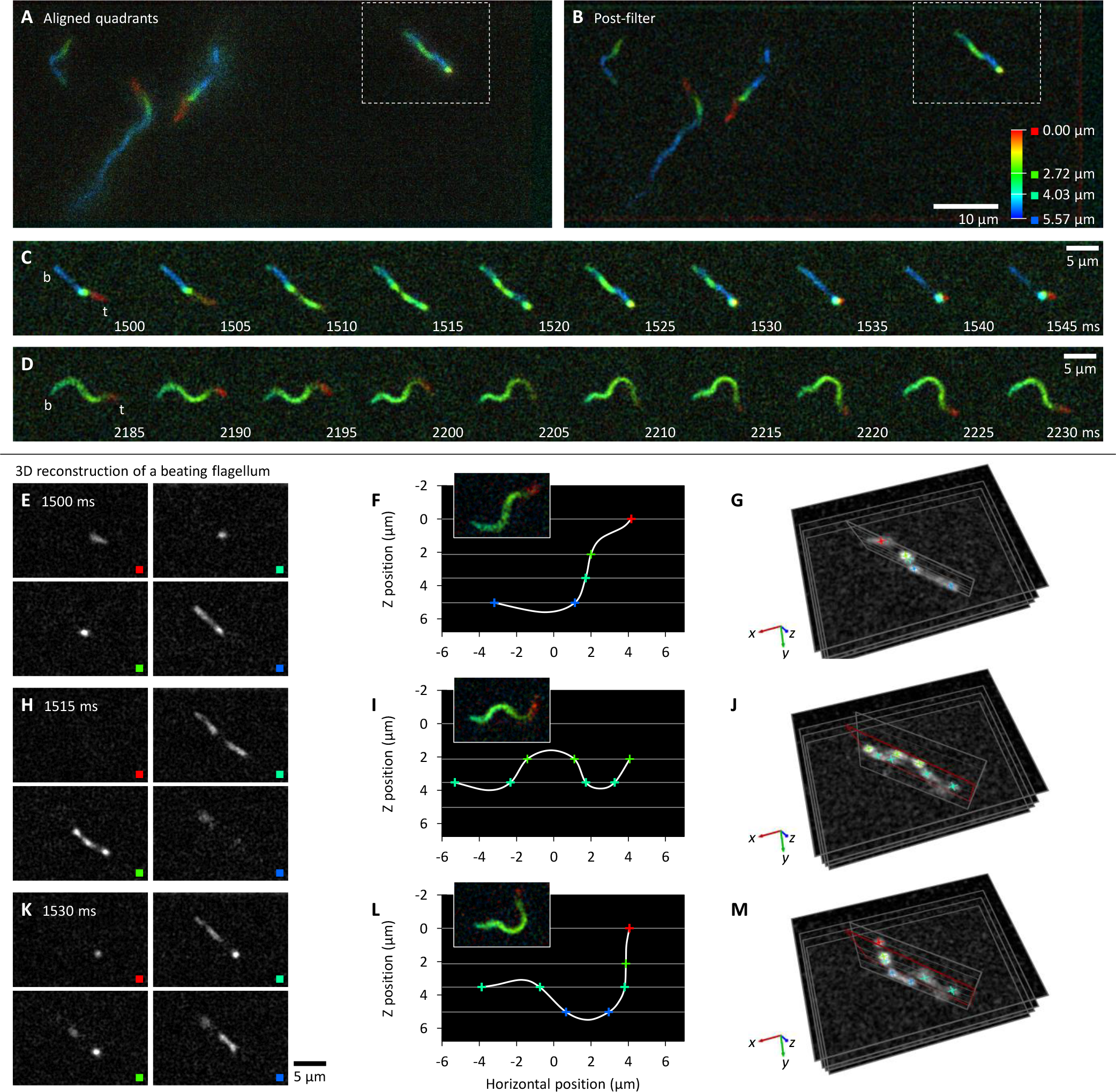
3D reconstruction of the *Leishmania mexicana* flagellar beat shows deviation from a plane. All panels show frames from a 200 Hz multifocal plane video of *L. mexicana* promastigotes expressing SMP1::mNG, a flagellar membrane protein. **A.** One example frame following alignment of the sub-images, pseudocoloured according to the depth scale. **B.** The image in A following filtering to aid interpretation. **C-D.** Movement of a single example flagellum, outlined in A and B, beating at ~20 Hz. b and t indicate the flagellum base and tip. **C.** One example beat cycle when the beat occurs perpendicular to the focal planes. **D.** One beat cycle from the same cell, but 0.7 s later when the cell has rotated such that the beat occurs parallel to the focal planes. **E-G.** 3D reconstruction of flagellum conformation from one frame (1530 ms in C) when beating perpendicular to the focal planes. **E.** Images for each focal plane. **F.** A plot of the points of intersection of the flagellum with the focal plane from the images in E. Inset image, a rotated copy of 2215 ms in D when the beat occurred parallel to the focal plane, for comparison. **F.** Direct visualisation of the focal planes in 3D. **H-J** and **K-M** are two further reconstructions of the same cell, at 1515 ms and 1530 ms respectively, as in E-G, showing the plane in G in red.

*Leishmania* were readily found with a flagellum undergoing a tip-to-base symmetrical flagellar beat with the beat plane oriented approximately parallel or perpendicular to the focal planes. A few cells rotated sufficiently quickly that, within the length of the video (3 s), both orientations could be seen for a single cell (Figure 3C,D). The 3D conformation of the flagellum could be reconstructed by recording the points where the flagellum intersected with a focal plane while beating perpendicular to the focal planes (Figure 3E-M). By considering a cell which rotates over the 3 s video, the conformation of the flagellum reconstructed from the multifocal plane imaging can be compared to the same flagellum at a similar stage of the flagellum beat once it has rotated parallel to the focal plane (Figure 3F,I,L). This confirmed an accurate 3D reconstruction.

The points of intersection of the flagellum with the focal plane can be accurately identified at high resolution as the computational alignment of the focal planes has a small error (Figure S1D). This approach was used to quantitatively determine flagellum beat planarity using frames of videos where the flagellum crossed the focal planes at 5 or more points. These points of were then fitted to a plane and the residuals (distance of a point from the fitted plane) used to give a measure of planarity. The mean residual indicates overall deviation from the plane, maximum residual gives an indication of maximum deviation from planarity at any single point. Overall, residuals were small (*n* = 23 reconstructions from 9 cells), indicating the configuration of the flagellum in any single frame is very close to planar (average mean residual 0.076±0.058 µm). Maximum residual was also small (average maximum residual 0.32±0.24 µm). This is <5% of average flagellum length^31^. However, while the flagellum configuration was near-planar in individual frames, this plane changed over the course of a beat, particularly for short flagella (illustrated in Figure 3C,D). As swimming is sensitive to subtle deviations of the flagellum beat from planar, as shown in sperm^22^, this deviation from planarity is likely to be important, perhaps for the characteristic helical paths of swimming *Leishmania*^2^.

### Conclusions

These analyses show it is feasible to use multifocal plane fluorescence microscopy to analyse volumes and frame rates useful for interrogating flagellum beating. It was successful with either chemical or native fluorescent protein fluorescence. This is among the highest frame rate biological applications of multifocal plane microscopy and is, to my knowledge, the first 3D reconstruction of flagellum-driven movement by fluorescence microscopy.

This approach is highly transferrable to other species or ciliated/flagellated tissue and has key advantages over transmitted light, as previously used in digital holography and bright-field defocus methods, coming from the specificity of fluorescent labels. This is powerful for direct analysis of flagellum beating, but there are also more complex applications, for example a labelled cell in a crowded environment, in tissue or among other cells – in trypanosomes such densely packed cells are where collective ‘social’ motility occurs^10^. Alternatively, the specific labelling of a cellular substructure is particularly powerful for complex multi-flagellated cells like *Giardia* and *Trichomonas*, but also specific flagellar or cellular structures in any organism. High-speed multifocal plane microscopy is therefore a powerful approach for understanding flagellum-driven motility.

## Methods

SmOxP9 procyclic *T. brucei*^32^ were grown in SDM79 with 10% FCS. Cas9T7 promastigote *L. mexicana*^33^ were grown in M199 (Life Technologies) supplemented with 2.2 g/L NaHCO_3_, 0.005% hemin, 40 mM Hepes⋅HCL (pH 7.4), and 10% FCS. *L. mexicana* SMP1 (LmxM.20.1310)^30^ was C-terminally tagged with mNG at the endogenous locus as previously described^33^. *T. brucei* and *L. mexicana* were grown at 28°C and imaged during logarithmic growth (0.5 to 1.0×10^7^ cells/ml).

Videos were captured using a 100× NA 1.4 objective (without phase ring, Zeiss) on an Axio observer A1 (Zeiss) microscope with incubator chamber, using a 120 V metal halide light source (Zeiss, HXP 120 V) and either an mRFP (Zeiss, 63HE) or GFP (ThorLabs, MDF-GFP2) filter cube. A MultiSplit v2 (Cairn Research) was used for multifocal plane imaging with three 50% semi-silvered mirrors in the filter cubes and a 2000 mm, 1300 mm, 500 mm or no lens in the auxiliary filter/lens mount immediately following the filter cube.

Images for calibration of focus offset, position offset and scale/magnification aberration of each sub-image were captured using multi-wavelength fluorescent beads (TetraSpeck, Invitrogen T7279). Focus offset was measured from the relative z position to focus the beads in each sub-image. Sub-image position offset and scale aberration were determined from bead pairs (one central and one peripheral bead within each sub-image) using Gaussian fitting to determine bead locations with sub-pixel accuracy.

Cells were held in a sealed 250 µm deep (deep relative to cell size) sample chamber (Gene Frame, ThermoFisher, AB0576) between a plain glass slide and coverslip in normal growth medium and analysed at 28°C. For FM 4-64FX (ThermoFisher Molecular Probes, F34653) labelling 1:100 40 µM was included in the medium.

475 ms videos at 200 Hz frame rate (5 ms/frame) were captured using a Neo 5.5 camera (Andor), vertically cropped to 50% (1080 px) around the camera midline. Constant readout column and pixel noise were subtracted (mean signal from 600 frames captured with no illumination) and random noise per row of pixels was subtracted (the median signal per pixel row per frame). Image stacks were then generated using the necessary offset and scale corrections. Image filtering for analysis was a 2 px Gaussian blur then a 10 px rolling ball background subtraction. Image processing was performed in ImageJ^34^.

## Acknowledgements

This work was supported by the Wellcome trust [211075/Z/18/Z, 104627/Z/14/Z, 103261/Z/13/Z]. I would like to thank Ziyin Wang for supplying the *L. mexicana* SMP1::mNG cell line and Keith Gull for his support through sharing lab equipment.

---

**Figure S1.**
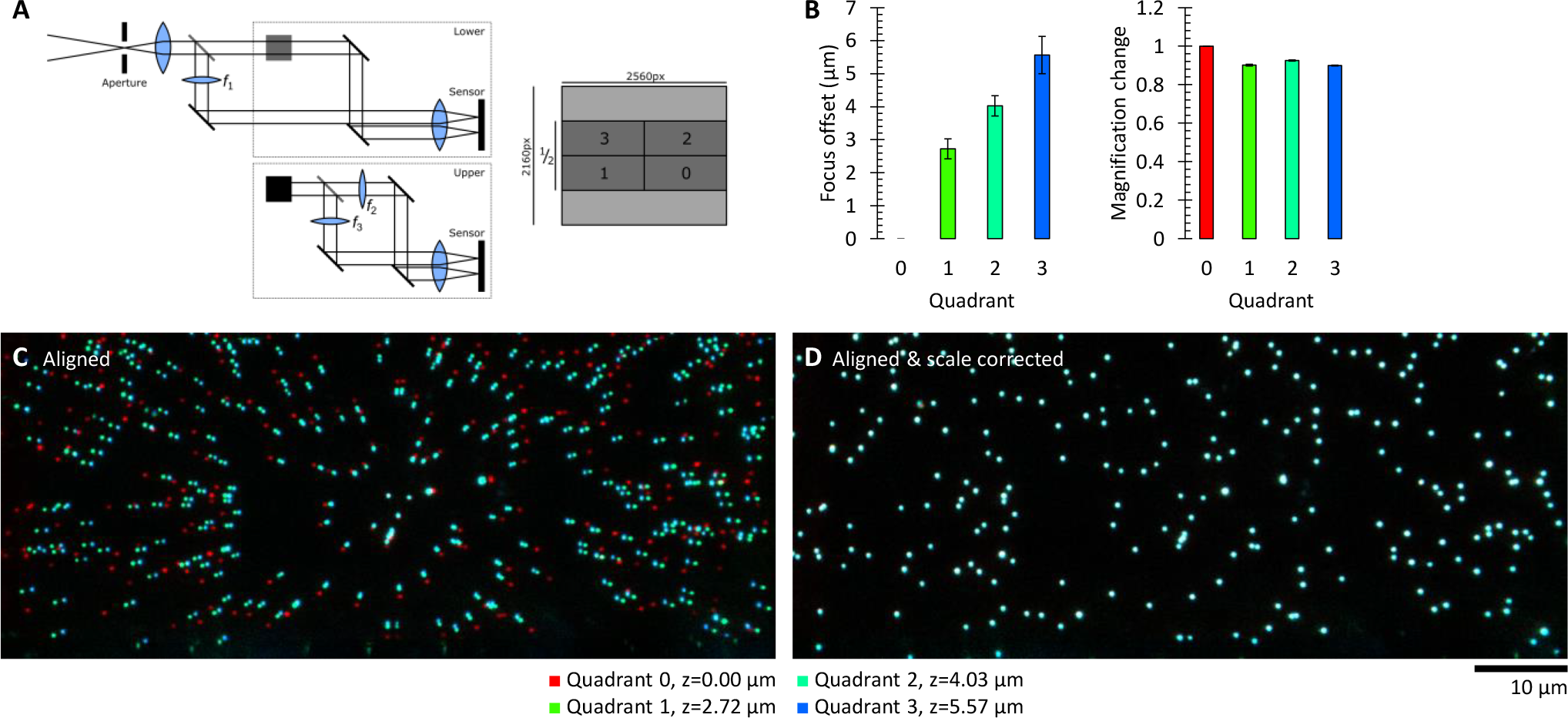
Setup and calibration of the multifocal plane system. **A.** Overview of the light path through the multi-splitter to achieve images of the same region but with offset focal planes on four quadrants of the camera sensor. Semi-silvered mirrors are shown in grey, fully-silvered in black. The light path is 3D, with the grey square (representing a semi-silvered mirror) in the lower section reflecting light upwards, out of the plane of the page/screen, and the black square (representing a fully-silvered mirror) in the upper section reflecting it back to parallel with the page/screen. Only half (full width, vertically centred) of the camera sensor was used, allowing a 200 Hz frame rate. *f*_1_ = 2000 mm, *f*_2_ = 1300 mm, *f*_3_ = 500 mm. **B.** The measured offset in focal plane for the four quadrants and the measured scale/magnification aberration for the four quadrants relative to the light path with no additional lens. Error bars represent the standard deviation, *n* = 5. **C.** Example of images aligned from the four quadrants (captured at the appropriate stage height for the beads to be in focus) without correction for the scale/magnification aberration. **D.** The same set of images aligned from the four quadrants with scale/magnification correction, showing precise co-localisation. Offset and scale parameters were accepted if the mean offset between points from different channels was <1 px (<65 nm).

